# IMMUNIA: A Multi-LLM Reasoning Agent for Immunoregulatory Surfaceome Discovery

**DOI:** 10.1101/2025.11.02.686138

**Authors:** Namu Park, Jung Hyun Lee

## Abstract

Biomarker discovery for immunotherapy often requires reasoning across complex immune contexts. We present IMMUNIA, a multi-large-language-model (multi-LLM) reasoning agent designed to identify immunoregulatory surfaceome genes through interpretable, biologically grounded analysis. The term IMMUNIA originates from the fusion of Immune and Noeia (the Greek concept of perception and understanding), defining an AI system that perceives, reasons, and interprets the immune landscape with human-like cognition. IMMUNIA integrates structured prompting, contextual scoring across immunotherapy, inflammation, and NF-κB signaling, and consensus reasoning across GPT-4o, GPT-5, and Gemini 2.5 Pro. Benchmarking with positive (HLA) and negative (contactin) controls confirmed model consistency and contextual discrimination. Consensus evaluation prioritized IL1R1, BSG, CD276, ALCAM, B2M, PTPRS, VCAN, and MXRA5 as high-confidence candidates. Among these, PTPRS, VCAN, and MXRA5 emerged as previously unrecognized stromal immune checkpoint-like regulators, shaping tumor-immune crosstalk via phosphatase, ECM, and cytokine signaling networks. IMMUNIA thus establishes a reasoning-centric AI paradigm that bridges computational inference with biological plausibility, offering a scalable approach for precision immunotherapy biomarker discovery.

## INTRODUCTION

Artificial intelligence has become indispensable in biomedical discovery, yet most current approaches remain constrained to pattern recognition and correlative prediction rather than reasoning about biological causality^1–3^. In immuno-oncology, where therapeutic outcomes depend on highly contextual interactions between tumor and immune cells, such correlation-based models often fail to capture the mechanistic underpinnings of response and resistance^4–6^. To translate the growing complexity of omics data into actionable biological understanding, next-generation AI systems must be capable of contextual reasoning, integrating mechanistic knowledge, cross-scale evidence, and interpretability^7^.

Cancer immunotherapy has transformed the therapeutic landscape across oncology, but its benefits remain unevenly distributed, with many patients exhibiting intrinsic resistance or relapse following initial response^8,9^. Overcoming these limitations requires biomarkers that not only correlate with clinical outcome but also illuminate the biological mechanisms that modulate anti-tumor immunity^10,11^. Among the most promising biomarker spaces is the immunoregulatory surfaceome, the ensemble of cell-surface and extracellular proteins that coordinate immune recognition, costimulation, and immune evasion^12–14^. This network includes well-characterized checkpoint molecules such as PD-1 and CTLA-4 as well as numerous underexplored regulators with potential to influence immune infiltration, antigen presentation, and effector function. Because of their accessibility to antibodies and biologics, these surface molecules represent a uniquely actionable frontier for precision immunotherapy^15,16^.

The expanding availability of bulk and single-cell transcriptomic, proteomic, and interactome datasets provides unprecedented opportunities to dissect the immunoregulatory surfaceome^17–19^. Yet translating these high-dimensional datasets into reproducible, clinically meaningful biomarkers remains challenging. Conventional bioinformatics pipelines, centered on differential expression and pathway enrichment, often overlook context-dependent regulators and lack mechanisms for integrating literature evidence, therapeutic data, or causal reasoning^20,21^. As a result, many potentially important immunomodulators remain hidden within existing datasets.

Large language models have recently emerged as powerful tools for synthesizing biomedical knowledge across unstructured text, databases, and experimental results^22,23^. Their ability to infer relationships and generate mechanistic hypotheses offers a new paradigm for data interpretation^24,25^. However, most applications still rely on single-model outputs and lack validation across independent reasoning systems. Integrating multiple large language models, each with distinct architectures, training corpora, and contextual reasoning strengths, can enhance robustness and reduce bias, similar to a panel of human experts providing complementary perspectives^26,27^. The rise of agentic AI frameworks, which coordinate autonomous model interactions and critical evaluation loops, further enables systematic and reproducible knowledge discovery^28^.

In this study, we develop and apply such an agentic multi model reasoning framework, termed IMMUNIA, to explore the immunoregulatory surfaceome and identify novel immunotherapy biomarkers. This approach aims to transform large-scale data interpretation from static analysis into a dynamic process of reasoning, consensus building, and mechanistic inference, providing a transparent and biologically grounded path toward precision immunotherapy discovery.

## METHODS

### Dataset acquisition and preprocessing

Bulk RNA-seq data from 84 prostate cancer patients were obtained from the Gene Expression Omnibus (accession GSE245708; Van Espen et al., 2023). Normalized expression matrices and metadata were directly used as provided in the public dataset. Gene identifiers were converted to current HGNC symbols, and duplicate entries were collapsed by median expression. Expression values were log1p-transformed for basic normalization. No additional preprocessing or quality control was applied beyond the procedures described in the original study.

### Gene curation and filtering

To restrict the analysis to extracellular immunoregulatory components, we curated immunoglobulin (Ig) domain–containing genes from InterPro and UniProt annotations, including Ig-like, V-set, I-set, C1-set, and C2-set subtypes. Redundant transcripts and pseudogenes were removed, resulting in 478 Ig-domain-containing surfaceome genes. Two gene subsets were generated: the top 10% ranked by (1) average expression and (2) expression variability across samples. The overlapping genes between these subsets were retained after excluding HLA-related entries, yielding 29 candidate genes used for subsequent IMMUNIA analyses.

### Prompt analysis

For each candidate gene, a concise functional context was assembled using GeneCards (function field only) and recent literature retrieved from PubMed describing immune, inflammatory, or signaling roles. This information was embedded in a standardized prompt that instructed the model to evaluate each gene’s relevance in three categories: Immunotherapy, Inflammation, and NF-κB signaling. Models assigned scores on a three-point scale, where 1 indicates low relevance, 2 moderate relevance, and 3 high relevance, and provided a brief rationale. Each response followed a strict JSON schema containing the gene symbol, three integer scores (in the order above), and a short rationale of no more than ten sentences. Responses not matching the schema were automatically reprompted with a format reminder and revalidated.

An example of the required output is as follows:

{ “Gene”: “PTPRS”, “Scores”: [1, 2, 3], “Rationale”: “PTPRS regulates the TLR9–NF-κB axis and cytokine output, indicating partial relevance to immunotherapy.” }

### Multi-LLM evaluation

Each structured prompt was independently analyzed by three large language models, GPT-4o, GPT-5, and Gemini 2.5 Pro, using a standardized schema that requested concise rationales and three category scores (Immunotherapy, Inflammation, and NF-κB signaling) on a one-to-three scale. From the curated set of 478 Ig-domain surfaceome genes, we focused on 29 prefiltered candidates selected based on combined expression level and variability criteria.

For each of these 29 genes, every model performed five independent reasoning runs with fixed random seeds, and the mean of the five runs per category was used as the model-specific score. Category scores were then summed across the three categories to obtain an overall relevance score, followed by ranking the genes according to their mean total scores. Genes consistently scoring highly across all three models were designated as overlapping high-confidence candidates for downstream analysis.

### Final candidate evaluation and downstream analysis

Top-ranked genes were reviewed and the overlapping high-confidence candidates across models were selected for downstream analysis. Each candidate’s model rationale was compared against GeneCards function information and PubMed literature to verify biological plausibility and novelty, removing genes with redundant or non specific reasoning and retaining those with clear, mechanistically supported explanations. Retained genes were annotated for therapeutic tractability by recording antibody accessibility, extracellular domain architecture, and any known drug or clinical trial associations. Biological context was mapped using Reactome style pathway groupings and protein interaction datasets to visualize relationships within broader immunoregulatory networks. Outputs were summarized as heatmaps of model derived scores, radar plots for the three relevance categories, and schematic diagrams depicting putative roles in the tumor immune microenvironment. Reproducibility was ensured by fixed model versions and version controlled prompts, schemas, and scripts, with JSON logs exported to preserve the full reasoning trail. The study used only publicly available, de identified transcriptomic data from GSE245708 and did not involve new human or animal experiments.

## Results

### IMMUNIA framework enables reasoning-centered biomarker discovery for precision immunotherapy

Immunotherapy has transformed cancer treatment, yet patient responses remain heterogeneous and difficult to predict. To address this challenge, we developed IMMUNIA, a reasoning-centered artificial intelligence framework that interprets immune regulatory mechanisms rather than relying solely on gene expression correlations. IMMUNIA integrates molecular data, literature evidence, and structured reasoning across multiple large language models to establish a transparent and reproducible process for identifying mechanistically relevant immunotherapy biomarkers.

As illustrated in Figure 1, the framework begins with biological relevance filtering of transcriptomic data from 84 prostate cancer patients (GEO accession GSE245708). From this dataset, we curated 478 Ig-domain–containing surfaceome genes, representing extracellular and plasma membrane proteins most likely to mediate immune recognition and therapeutic accessibility. These genes defined the candidate search space for downstream reasoning analysis.

**Figure 1.**
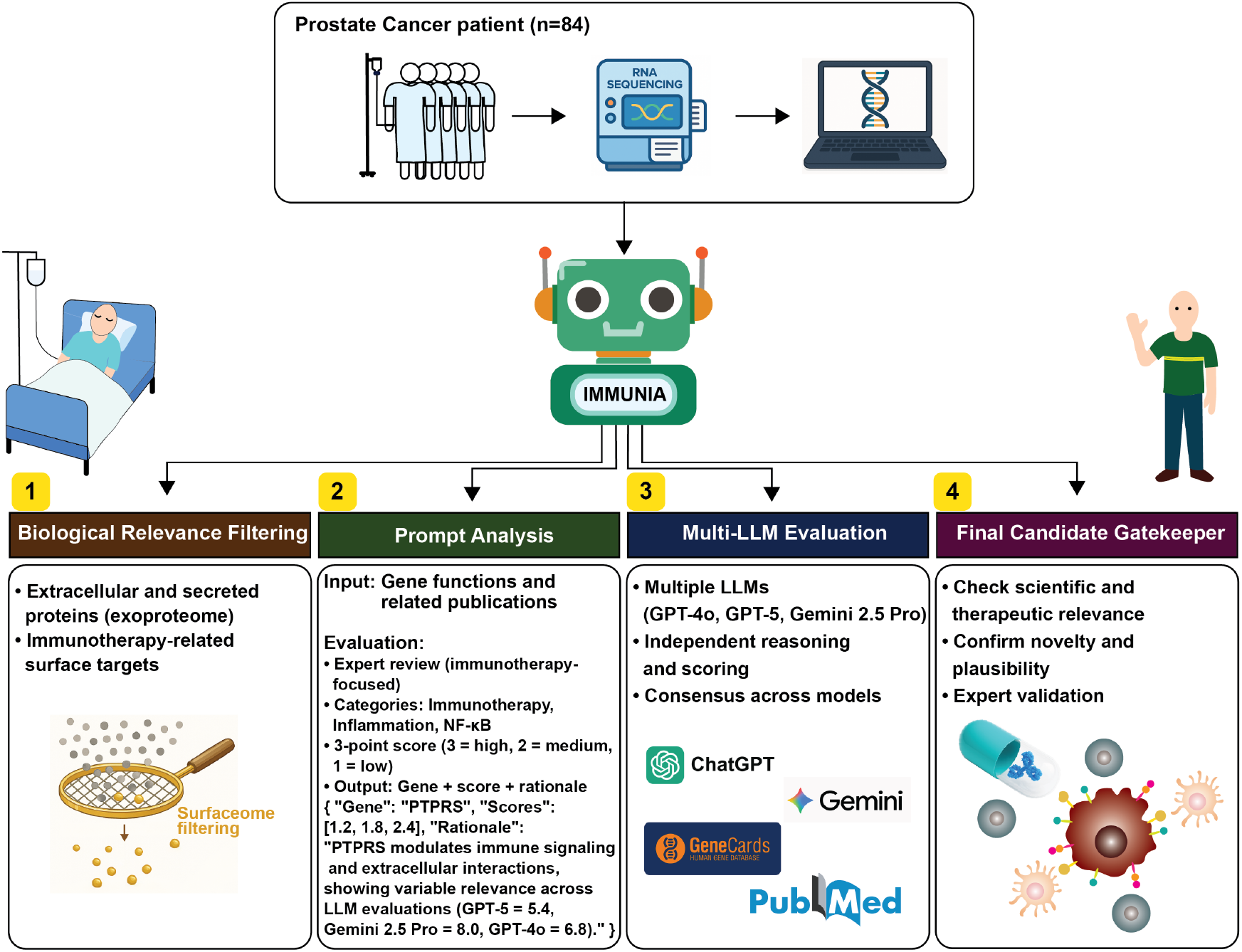
Multi-LLM reasoning framework for immunoregulatory biomarker discovery. Workflow illustrating the four-step analytical process applied to prostate cancer RNA-seq data (GSE245708, n = 84). Candidate surfaceome genes (n = 478) were filtered for extracellular and Ig-domain proteins (Step 1), analyzed using structured prompts integrating GeneCards and PubMed data (Step 2), independently evaluated across GPT-4o, GPT-5, and Gemini 2.5 Pro with five-run averaging per model (Step 3), and finalized through consensus-based expert validation for novelty and therapeutic relevance (Step 4).

In the prompt analysis stage, each gene was characterized using concise functional summaries from GeneCards and supporting literature from PubMed describing immune, inflammatory, or NF-κB– related functions. These elements were incorporated into standardized prompts that instructed models to assign three relevance scores (1 = low, 2 = moderate, 3 = high) for Immunotherapy, Inflammation, and NF-κB signaling, along with a brief textual rationale.

The multi-LLM evaluation stage employed GPT-4o, GPT-5, and Gemini 2.5 Pro, which independently analyzed the 29 prefiltered candidate genes using identical prompt structures. Each model performed five independent reasoning runs per gene with fixed random seeds to minimize stochastic variation. The mean score across runs and categories was used as the final model-specific value, and results were aggregated into a structured reasoning matrix.

Finally, in the candidate gatekeeper stage, genes were ranked by their mean cross-model relevance scores, and overlapping high-confidence candidates were identified. Each rationale was reviewed against GeneCards and PubMed to confirm biological plausibility and highlight potential novelty.

By combining reproducible multi-model reasoning with expert-style evidence synthesis, IMMUNIA converts transcriptomic data into interpretable, mechanistically grounded biomarker insights. This framework offers a scalable strategy for discovering immunoregulatory surface targets that link gene-level mechanisms to therapeutic potential in precision immunotherapy.

### Structured prompt design for cancer type-specific biomarker discovery

To ensure that IMMUNIA reasoning outputs could be generalized beyond a single tumor type, we implemented a structured prompt framework that defines context-specific evaluation categories for different cancer systems (Figure 2). In the present study, all analyses were performed using the general cancer immunotherapy context, which evaluates genes across three axes: immunotherapy relevance, inflammatory regulation, and NF kappa B signaling. These dimensions were selected because they capture core biological processes that govern antitumor immune activity, checkpoint responsiveness, and cytokine-driven signaling. This general framework provides a scalable baseline that can be extended to disease-specific prompts for precision biomarker discovery.

**Figure 2.**
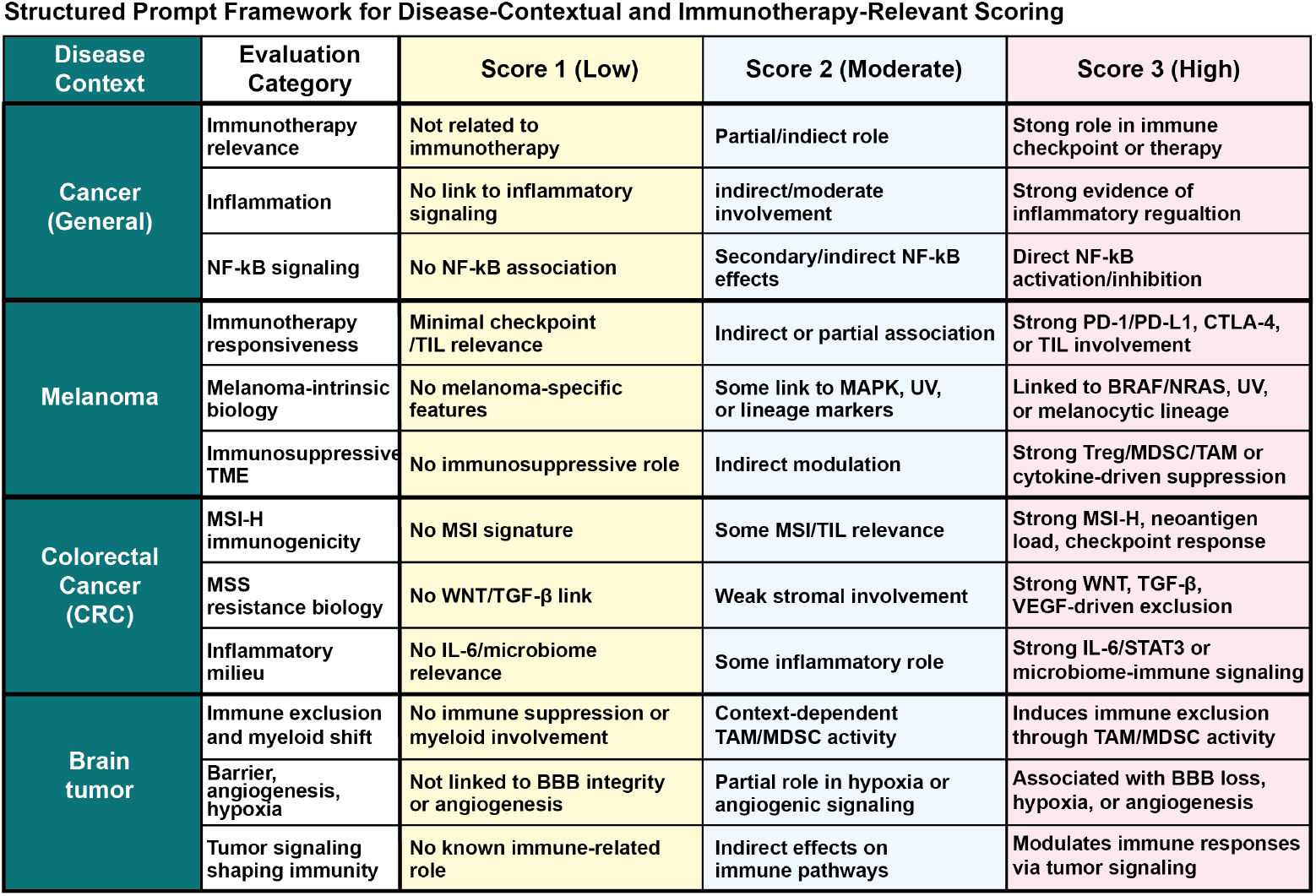
Structured prompt framework for disease contextual and immunotherapy relevant scoring. The scoring framework defines standardized evaluation criteria used by large language models to assess gene relevance across immune and inflammatory contexts. Each gene is scored on a three point scale (1 = low, 2 = moderate, 3 = high) according to its reported or inferred relationship to immunotherapy, inflammation, and NF kappa B signaling. The “Cancer (General)” section summarizes the categories applied in this study, while additional contexts including melanoma, colorectal cancer (CRC), and brain tumor illustrate extended disease specific templates for future applications. This schema enables consistent and interpretable scoring of gene immune relationships across diverse tumor types.

To demonstrate the adaptability of the framework, additional prompt templates were developed for melanoma, colorectal cancer (CRC), and brain tumors. Each cancer type required modifications that reflect unique features of its tumor immune microenvironment and molecular pathophysiology.

For melanoma, the evaluation axes included (1) immunotherapy responsiveness, focusing on PD-1 and CTLA-4 checkpoint sensitivity; (2) melanoma-intrinsic biology, encompassing MAPK, UV, and lineage-associated markers; and (3) immunosuppressive tumor microenvironment, emphasizing Treg, MDSC, and TAM-mediated suppression.

For colorectal cancer, the categories addressed (1) MSI-H immunogenicity, linked to microsatellite instability and neoantigen burden; (2) MSS resistance biology, characterized by WNT, TGF beta, and stromal exclusion pathways; and (3) inflammatory milieu, reflecting IL-6 and microbiome-associated signaling.

For brain tumors, prompts focused on (1) immune exclusion and myeloid shift, involving TAM and MDSC dynamics; (2) barrier, angiogenesis, and hypoxia signatures, representing BBB integrity and metabolic adaptation; and (3) tumor signaling shaping immunity, addressing pathways that indirectly regulate immune infiltration.

Each disease-specific schema retains the same three-level scoring scale (1 = low, 2 = moderate, 3 = high relevance) but applies it to cancer-specific biological contexts. This modular structure allows IMMUNIA to be rapidly adapted to diverse malignancies while maintaining a consistent reasoning format. By combining standardized scoring with contextual definitions, the framework supports systematic biomarker prioritization not only in general immunotherapy discovery but also within specialized tumor ecosystems that exhibit distinct immunologic signatures.

### Validation using positive and negative control genes in immunotherapy context

To evaluate the reliability of reasoning-based scoring, we performed an internal validation using Gemini 2.5 Pro, applying the IMMUNIA prompt schema specifically from the immunotherapy perspective. Because reasoning-based biomarker discovery is still an emerging approach, there are currently no standardized definitions of “positive” or “negative” control genes for AI-based immunotherapy evaluation. Nevertheless, incorporating such reference sets provides an essential benchmark for assessing biological plausibility and ensuring interpretive consistency across models.

As shown in Figure 3, we selected representative genes from two distinct categories within the surfaceome. The positive control group included canonical immune regulatory genes from the HLA family (HLA-A, -B, -C, -F, and -G), which are well known for their roles in antigen presentation and immune recognition. In contrast, the negative control group consisted of surface adhesion molecules (CNTN3–CNTN6, OPCML) that are structurally similar but lack recognized immune-modulatory functions.

**Figure 3.**
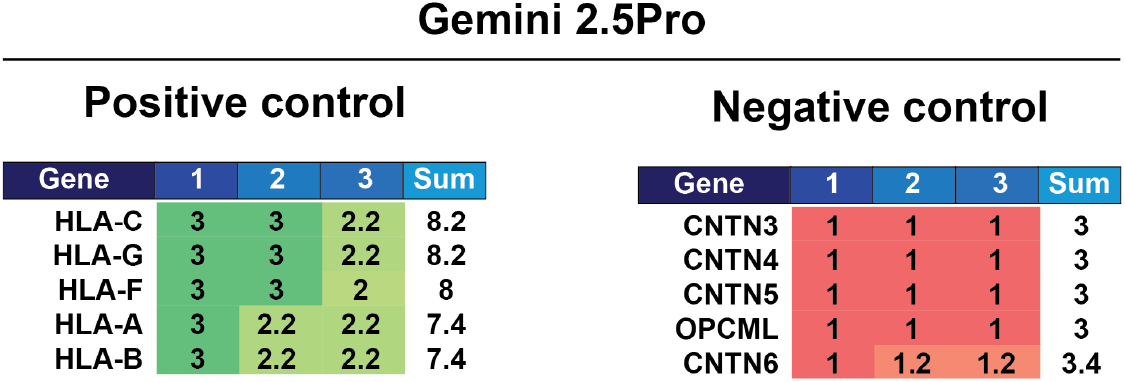
Positive and negative control validation using Gemini 2.5 Pro. To assess scoring reliability and contextual reasoning, Gemini 2.5 Pro was tested with known immune positive and negative control genes. Positive controls (HLA class I and non-classical HLA genes) consistently received high relevance scores across immunotherapy, inflammation, and NF kappa B categories, reflecting their well-established immune regulatory roles. In contrast, negative controls (contactin and adhesion-related genes) uniformly scored low, indicating no immunologic association. These results demonstrate the model’s ability to distinguish immune active from non-immune genes within the surfaceome, confirming interpretive stability of the structured prompt framework.

Gemini 2.5 Pro produced a clear and biologically coherent separation between these two sets. HLA genes consistently received high cumulative scores (7.4–8.2) across the three evaluation axes (immunotherapy relevance, inflammation, and NF kappa B signaling), while contactin and OPCML genes scored uniformly low (around 3.0–3.4), reflecting an absence of immunologic reasoning in model outputs. These results demonstrate that IMMUNIA’s reasoning-based evaluation, even within a single LLM, can recapitulate biologically expected patterns of immune relevance among surfaceome genes.

Although the concept of positive and negative controls has not yet been formalized in AI-driven immunotherapy research, the ability to distinguish between established immune regulators and nonimmune structural proteins provides an important internal reference point. Incorporating such validation steps will be critical for defining biologically meaningful boundaries in future reasoning-based biomarker discovery studies.

### Cross model comparison of reasoning patterns across 29 immunoregulatory surface genes

To evaluate the robustness and reproducibility of multi model reasoning, three large language models, GPT 4o, GPT 5, and Gemini 2.5 Pro, were compared using a curated panel of 29 immunoregulatory surfaceome genes (Figure 4). Each model independently analyzed all genes through five separate reasoning runs, and the mean of these runs was used to calculate the final score for each of the three evaluation axes: immunotherapy relevance, inflammation, and NF kappa B signaling.

**Figure 4.**
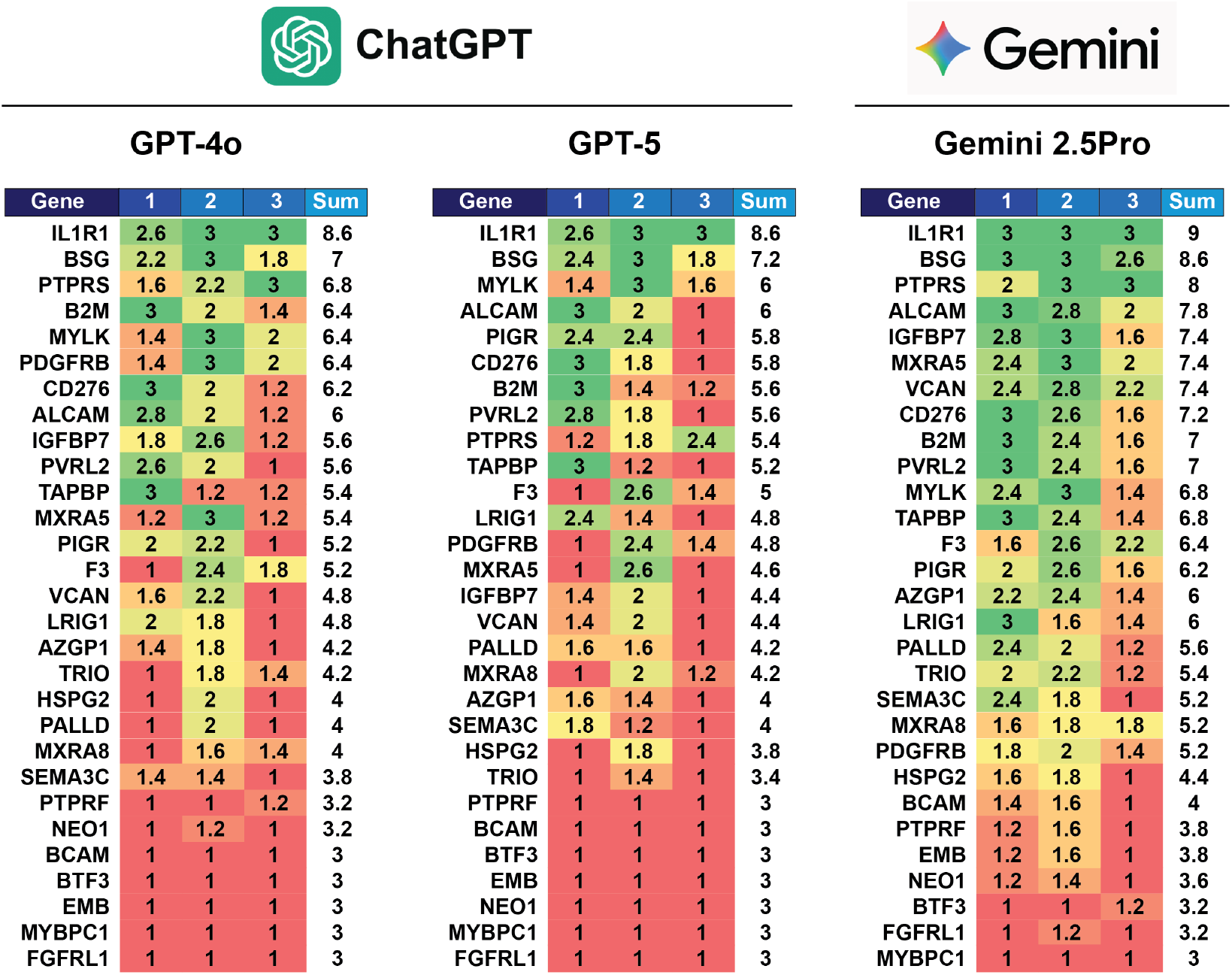
Cross-model comparison of gene-level relevance scoring by GPT-4o, GPT-5, and Gemini 2.5 Pro. Heatmaps show averaged scores from five independent reasoning runs for 29 surfaceome genes evaluated across three large language models. Each gene was scored for immunotherapy, inflammation, and NF kappa B relevance on a three point scale, with the total sum reflecting overall immunologic relevance. High scoring genes such as IL1R1, BSG, and PTPRS were consistently prioritized across models, indicating strong cross-model agreement. FGFRL1 and MYBPC1 scored lowest in all models, serving as internal negative references. While GPT-4o emphasized mechanistic reasoning, GPT-5 expanded causal interpretation, and Gemini 2.5 Pro provided broader system-level context, together demonstrating reproducible yet complementary reasoning patterns across architectures.

Across all models, the overall scoring patterns were highly consistent, indicating stable contextual reasoning. Genes such as IL1R1, BSG, and PTPRS repeatedly ranked among the highest scoring candidates, consistent with their known or predicted involvement in immune regulation and tumor immune interactions. Similarly, ALCAM, B2M, and CD276 maintained intermediate to high scores across the three models, aligning with their reported functions in antigen presentation, stromal signaling, and checkpoint regulation. This pattern demonstrates that the structured evaluation schema enables each model to independently converge on biologically coherent interpretations.

Distinct reasoning styles emerged among models. GPT 4o favored mechanistic explanations, assigning higher scores to genes linked to receptor ligand signaling or cytokine activity. GPT 5 provided more conservative but causally integrated assessments, emphasizing pathway level consistency and penalizing genes with limited experimental support. Gemini 2.5 Pro produced broader contextual reasoning, frequently incorporating microenvironmental and inflammatory feedback processes, which resulted in slightly higher overall averages and a smoother scoring distribution across categories.

Conversely, several genes consistently received very low scores in all three models, including FGFRL1 and MYBPC1, both of which lack known immunoregulatory or inflammatory functions. Their uniformly low ranking across five independent runs per model highlights the specificity of the reasoning based evaluation, distinguishing biologically irrelevant structural or metabolic genes from those with immune related potential.

Together, these observations confirm that independent large language models, despite their architectural and reasoning differences, can reproducibly identify common immunologic patterns while maintaining discrimination against unrelated candidates. The convergence of high scoring immunoregulatory genes and uniformly low scoring negatives supports the reliability and interpretive precision of multi model reasoning for immune relevant biomarker evaluation.

### Consensus prioritization and therapeutic landscape of high scoring genes

The cross model evaluation of the 478 Ig domain surfaceome genes yielded a common top ten set relevant to immunotherapy: IL1R1, BSG, CD276, ALCAM, B2M, PTPRS, VCAN, MXRA5, F3, and PVRL2 (Figure 5). We summarized the current therapeutic status for each target using public sources. CD276 has multiple antibodies and antibody drug conjugates in clinical trials. F3 has an approved antibody drug conjugate for cervical cancer. PVRL2 has agents in early clinical development. BSG has a humanized antibody evaluated in exploratory studies. IL1R1 has neutralizing antibodies at preclinical or research stages while approved drugs in this pathway currently target IL 1β or IL 1RA. ALCAM and B2M have experimental reagents reported but no approved therapy.

**Figure 5.**
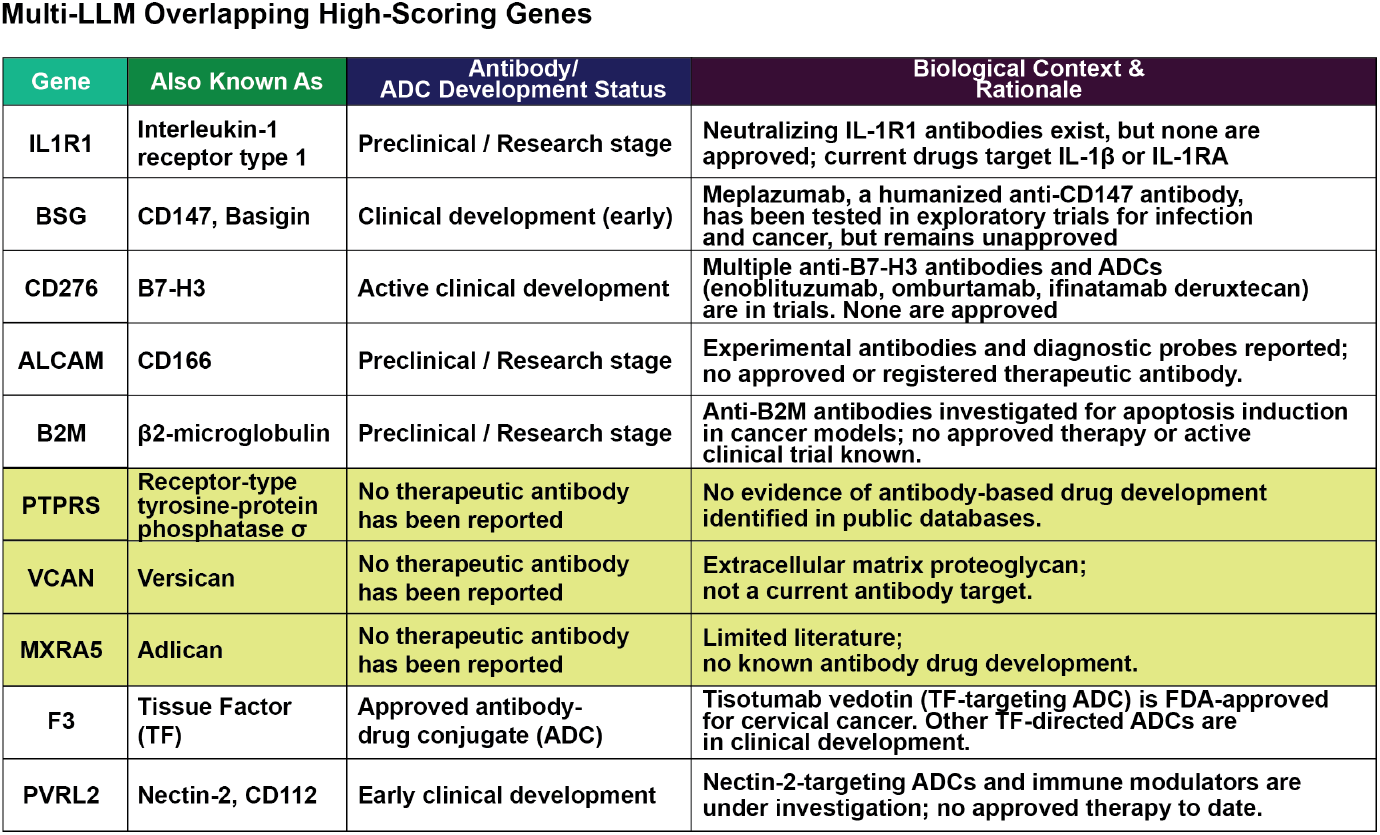
Summary of high scoring immunoregulatory surfaceome genes across multiple LLMs. The ten highest scoring genes identified through consensus among GPT-4o, GPT-5, and Gemini 2.5 Pro are shown with their corresponding therapeutic development status and biological rationale. Well-known immunotherapy targets such as IL1R1, CD147 (BSG), and B7-H3 (CD276) are already under preclinical or clinical evaluation, validating the model’s predictive consistency. In contrast, PTPRS, VCAN, and MXRA5 have no reported antibody or ADC development, indicating novel immunoregulatory potential. These genes are associated with extracellular matrix remodeling, phosphatase signaling, and stromal immune modulation, suggesting roles distinct from canonical immune checkpoints. The integration of reasoning outputs and therapeutic databases highlights unmet opportunities for developing next-generation immunomodulatory antibodies targeting the tumor microenvironment.

In contrast, PTPRS, VCAN, and MXRA5 show no evidence of ongoing therapeutic antibody development despite high and consistent relevance scores from all three language models. We therefore selected these three genes for deeper biological interrogation in subsequent figures.

Placing these results in context, monoclonal antibodies against PD 1 and PD L1 established modern immunotherapy by demonstrating that extracellular immune regulators can be drugged with high specificity to restore antitumor T cell activity. Success with PD 1 and PD L1 illustrates three principles that guide target nomination. First, targets must sit at critical control points of immune recognition or inflammatory signaling. Second, targets should be extracellular or cell surface to enable pharmacologic access by antibodies and biologics. Third, targets need a mechanistic rationale that links modulation to measurable immune outcomes such as antigen presentation, myeloid polarization, or cytokine balance.

The top ten list satisfies the second criterion by design, and several members satisfy the first through established roles in antigen presentation or checkpoint signaling. The absence of active drug programs for PTPRS, VCAN, and MXRA5 highlights an actionable gap. Their extracellular localization and model derived rationales that implicate phosphatase control of NF kappa B and STAT signaling, matrix mediated regulation of cytokine diffusion and myeloid recruitment, and stromal remodeling associated with immune suppression indicate credible mechanisms that parallel the logic of successful checkpoint targets. These observations justify focused follow up that includes epitope mapping, antibody generation, and functional assays in immune co culture systems to determine whether modulating these proteins can reshape the tumor immune environment and restore antitumor immune activity.

### Comparative reasoning analysis enhances biomarker interpretability

To examine how large language models differ in their biological reasoning and to assess the interpretive reliability of their outputs, we performed a focused analysis on PTPRS, one of the top scoring but previously underexplored genes identified in the surfaceome screen (Figure 6). PTPRS encodes a receptor type tyrosine phosphatase with potential roles in immune regulation through NF kappa B and cytokine signaling. Each model, GPT 4o, GPT 5, and Gemini 2.5 Pro, analyzed the gene independently using the same structured prompt, scoring system, and reasoning schema. The results reveal not only consistent biological logic across models but also distinct reasoning styles that together expand interpretive depth.

**Figure 6.**
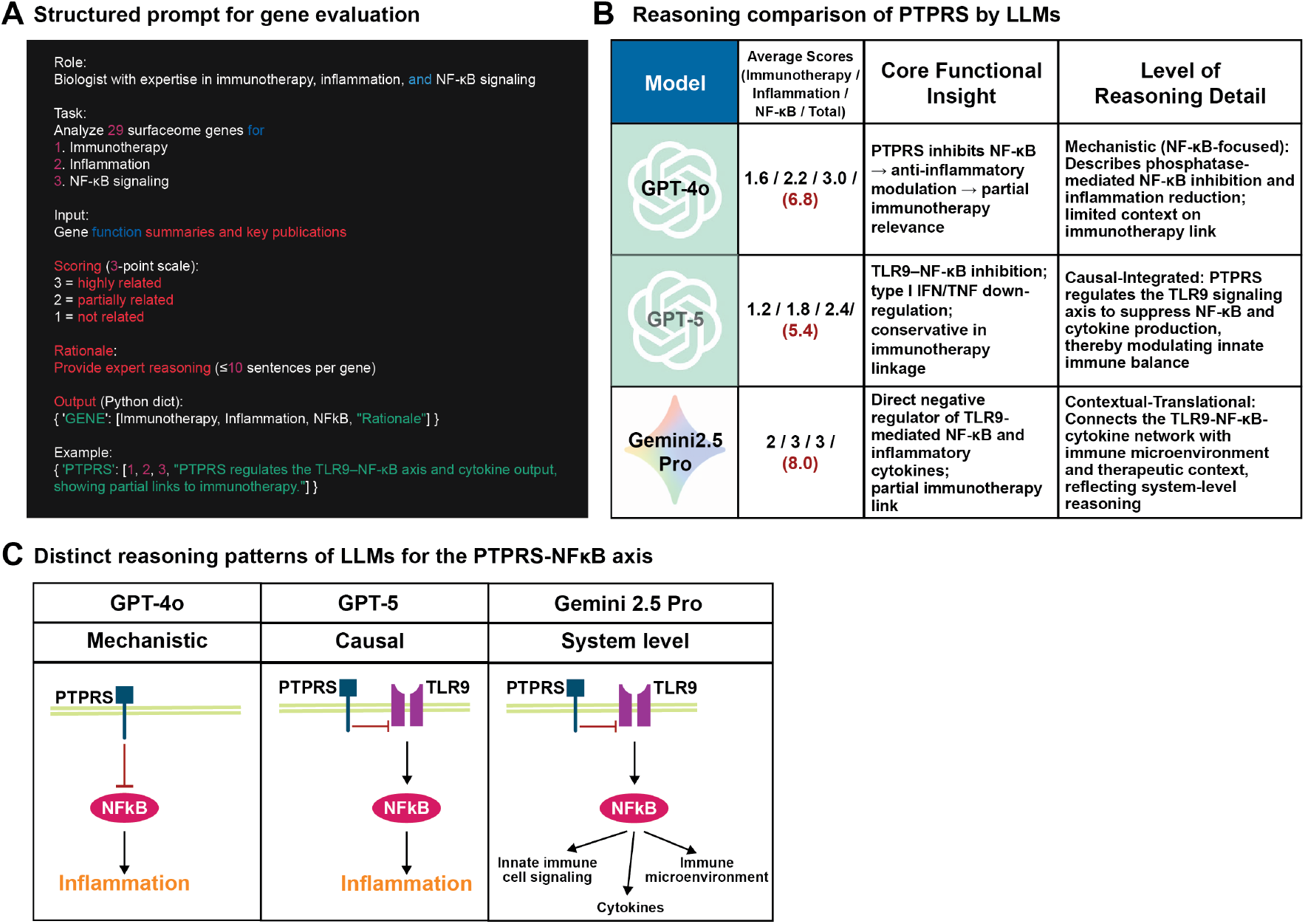
Comparative reasoning analysis of PTPRS across multiple large language models. (A) Structured prompt used for systematic evaluation of 29 surfaceome genes. Each gene was analyzed for immunotherapy relevance, inflammation, and NF kappa B signaling using a three point scale, accompanied by concise textual reasoning. (B) Comparison of reasoning outputs for PTPRS across GPT-4o, GPT-5, and Gemini 2.5 Pro. GPT-4o emphasized mechanistic phosphatase-mediated NF kappa B inhibition, GPT-5 described causal regulation through the TLR9 NF kappa B signaling axis, and Gemini 2.5 Pro expanded to system-level reasoning linking cytokine networks and the tumor microenvironment. (C) Conceptual schematic illustrating model-specific reasoning depth, from molecular mechanism to causal integration and contextual immune modulation. This multi-model reasoning triangulation enables robust hypothesis generation for gene-level immune function analysis.

As shown in Figure 6A, the standardized prompt required each model to evaluate gene relevance to immunotherapy, inflammation, and NF kappa B signaling, assigning three point scores (1 low, 2 moderate, 3 high) and providing a concise expert style rationale. The outputs were then compared quantitatively and qualitatively. In Figure 6B, GPT 4o emphasized mechanistic reasoning, describing phosphatase mediated inhibition of NF kappa B and its link to inflammation control, while providing limited discussion of immunotherapy relevance. GPT 5 generated causal integrated reasoning, situating PTPRS within the TLR9 NF kappa B signaling axis and suggesting that it modulates cytokine production to influence innate immune balance. Gemini 2.5 Pro exhibited contextual translational reasoning, connecting the phosphatase’s function to cytokine networks, immune microenvironment feedback, and therapeutic implications, reflecting a system level understanding.

The schematic summary in Figure 6C illustrates this gradient of interpretive complexity. GPT 4o focuses on local molecular interactions, GPT 5 extends to pathway level causality, and Gemini 2.5 Pro incorporates microenvironmental context. Together, these complementary perspectives demonstrate how multi model reasoning enhances biomarker discovery by integrating molecular mechanisms with immunologic systems level insight.

Such comparative reasoning is a critical step for improving biomarker accuracy and reproducibility. It allows validation not only of numerical scoring consistency but also of the underlying scientific logic guiding model outputs. In practical terms, this process functions as an internal interpretive checkpoint, ensuring that candidate genes such as PTPRS are supported by coherent biological rationale before advancing to experimental validation or therapeutic exploration.

### Identification of novel immunoregulatory surfaceome genes shaping the tumor–immune interface

The final integrative analysis highlighted PTPRS, VCAN, and MXRA5 as previously unrecognized immunoregulatory surfaceome genes with strong contextual links to immune modulation (Figure 7). Each gene represents a distinct but complementary mechanism of immune control within the tumor microenvironment. PTPRS, a receptor-type phosphatase, emerged as a potential intracellular immune checkpoint–like modulator that regulates cytokine and STAT3 signaling to fine-tune NF kappa B–mediated inflammation and immune homeostasis. VCAN, an extracellular matrix proteoglycan, was inferred to remodel the tumor stroma by limiting T cell infiltration and promoting M2 macrophage polarization through NF kappa B and TGF beta signaling. MXRA5, a secreted glycoprotein with multiple Ig like domains, was associated with PD L1 and B7 H3 expression, suggesting a stromal checkpoint like function that facilitates immune evasion in mesenchymal tumors.

**Figure 7.**
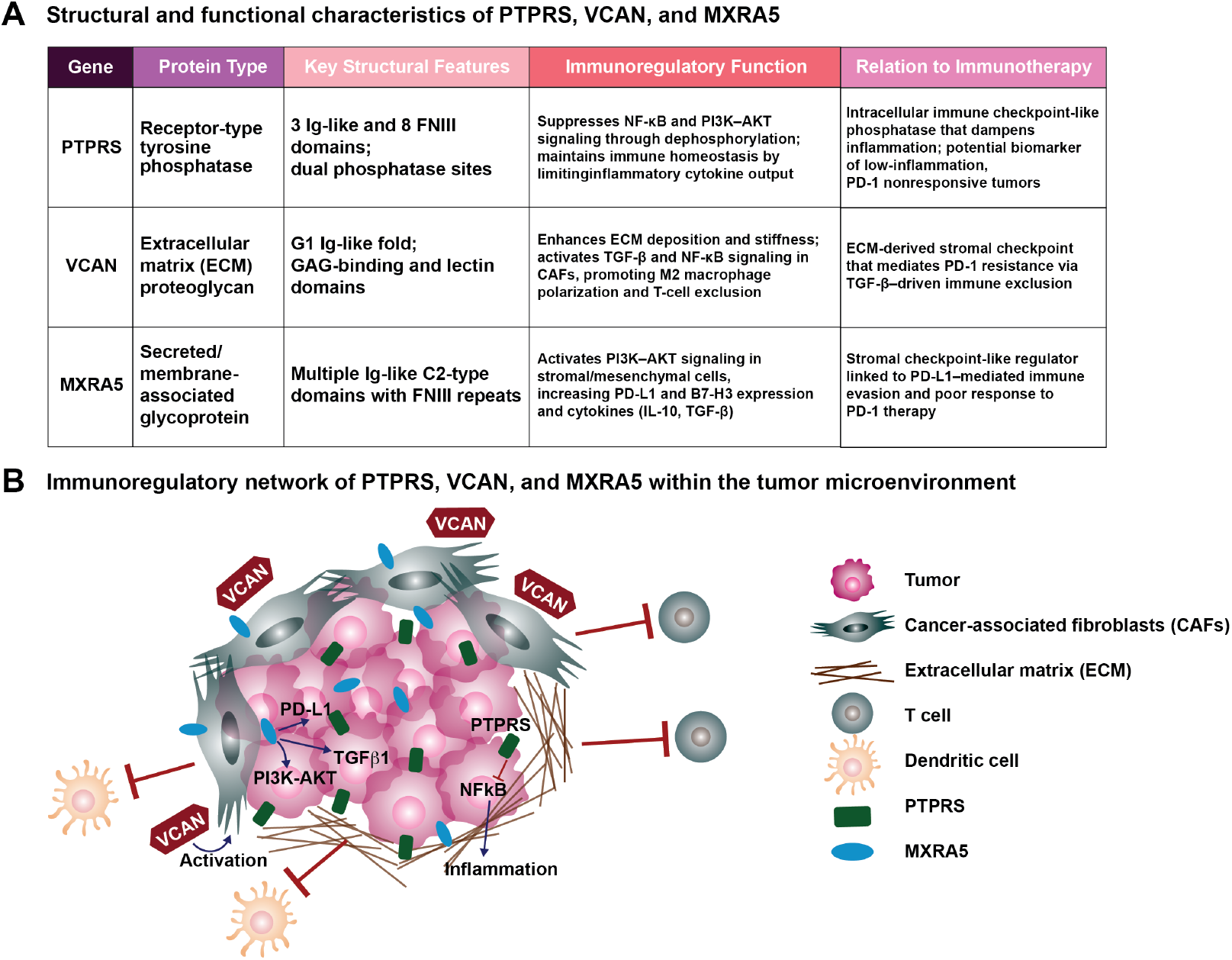
Identification and functional mapping of previously unrecognized immunoregulatory surfaceome genes. (A) Summary of three high-confidence candidates, PTPRS, VCAN, and MXRA5, identified through multi-LLM consensus as novel immunoregulatory surfaceome genes. Each exhibits distinct structural and functional features that suggest regulatory roles at the tumor–immune interface. PTPRS functions as a phosphatase controlling STAT3 and NF kappa B signaling, VCAN acts as an extracellular matrix proteoglycan modulating cytokine diffusion and T cell infiltration, and MXRA5 is a secreted glycoprotein linked to immune suppressive stromal remodeling. (B) Conceptual model illustrating how these molecules interact within the tumor microenvironment. PTPRS may suppress inflammatory NF kappa B signaling, VCAN forms a matrix barrier influencing cytokine gradients and immune cell accessibility, and MXRA5 supports immune suppressive fibroblast and macrophage activity. Together, they define a new class of stromal checkpoint-like regulators that shape immune evasion and therapeutic response in solid tumors.

The schematic model in Figure 7B visualizes how these molecules cooperate within the tumor and its surrounding microenvironment. PTPRS may regulate immune activation thresholds through phosphatase control of NF kappa B signaling, VCAN acts as a cytokine buffering scaffold that restricts immune cell access, and MXRA5 mediates stromal signaling feedback that reinforces macrophage driven immune suppression. Together, they illustrate a broader concept of stromal immune checkpoints, noncanonical regulators that shape antitumor immunity not through direct ligand receptor blockade but through biochemical and structural modulation of the tumor milieu.

These findings underscore the power of multi LLM reasoning to move beyond correlation based biomarker discovery toward mechanistically interpretable predictions. By integrating functional reasoning with biological plausibility, this study reveals an uncharted layer of immunoregulation at the tumor stroma interface. The discovery of PTPRS, VCAN, and MXRA5 not only expands the catalog of potential immunotherapy targets but also opens a new conceptual frontier in precision immuno oncology, where AI guided reasoning can illuminate how the immune system perceives, interprets, and responds to cancer.

## Discussion

This study establishes a reasoning centered framework for biomarker discovery in cancer immunotherapy by coordinating structured prompting, independent analysis, and consensus synthesis across multiple large language models. Through this approach, complex biological information is interpreted not as isolated data points but as interconnected narratives linking gene expression, molecular structure, and immune function. This design enables systematic identification of immunoregulatory surface genes that operate beyond canonical immune checkpoints and provides a transparent logic trail connecting computational inference to biological plausibility.

The comparative evaluation across GPT 4o, GPT 5, and Gemini 2.5 Pro demonstrated that different models contribute distinct layers of reasoning that collectively improve interpretive depth. GPT 4o emphasized molecular mechanisms such as receptor signaling and cytokine regulation, GPT 5 produced causal reasoning that connected pathway dynamics with immune activation, and Gemini 2.5 Pro delivered system level reasoning by integrating tumor microenvironmental context and translational relevance. The convergence of these complementary outputs across five independent runs per model produced stable, reproducible results, analogous to multi expert consensus in scientific peer evaluation.

Within this framework, three previously unrecognized immunoregulatory surface genes, PTPRS, VCAN, and MXRA5, consistently emerged as high confidence candidates. All three genes reside at the intersection of tumor, stroma, and immune cells, a regulatory niche that strongly influences cytokine gradients and immune accessibility yet remains underrepresented in current immunotherapy research. PTPRS, a receptor type phosphatase, was associated with STAT3 and NF kappa B regulation and linked to macrophage polarization and inflammation resolution. VCAN, an extracellular matrix proteoglycan, was repeatedly interpreted as a modulator of cytokine diffusion and leukocyte infiltration. MXRA5, a secreted matrix glycoprotein, appeared to reinforce an immunosuppressive microenvironment through interactions related to PD L1 and B7 H3 signaling. Together, these findings highlight a class of stromal immune checkpoint like regulators that may control immune activity through matrix remodeling and paracrine signaling rather than direct ligand receptor inhibition.

The therapeutic landscape of the top ten high scoring genes reveals a clear translational gap. Among them, only F3 (Tissue Factor) currently has an FDA approved agent, the antibody drug conjugate tisotumab vedotin (Tivdak) for cervical cancer. Other known regulators such as CD276 (B7 H3), BSG (CD147), and PVRL2 (Nectin 2) remain in early to mid stage clinical trials, while PTPRS, VCAN, and MXRA5 have no reported therapeutic programs. Their consistent prioritization despite the absence of prior drug development underscores their novelty and biological potential as next generation immunomodulatory targets.

Future studies should aim to elucidate how these candidates function at the protein interaction level. Detailed investigation into extracellular binding partners, matrix interactions, and effects on immune cell activation or infiltration will be critical for understanding their roles within the tumor microenvironment. For example, identifying whether PTPRS engages specific cytokine receptors or adhesion molecules could clarify its impact on macrophage or T cell signaling. Defining how VCAN organizes proteoglycan networks that bind chemokines or growth factors may reveal its control over leukocyte migration, while mapping MXRA5 interactions could explain its role in stromal feedback and myeloid activation. These studies will determine whether modulation of these molecules can reprogram immune activity and provide the foundation for developing therapeutic antibodies or biologics against them.

Methodologically, this work demonstrates that coordinated reasoning among multiple large language models can overcome the bias of single model prediction and introduce transparency into biological interpretation. Each reasoning step is documented in a machine readable format, allowing full traceability from prompt to conclusion. This auditability provides a tangible implementation of explainable artificial intelligence in biomedicine, converting qualitative reasoning into structured evidence that can be cross validated by human experts. As multi model reasoning matures, its application could extend to related areas such as drug mechanism elucidation, pathway curation, and therapeutic repurposing.

In conclusion, this study proposes a scalable and interpretable paradigm for AI driven biomarker discovery that integrates computational reasoning with biological insight. By combining multi model consensus with mechanistic interpretation, we identified both established and novel immune regulators, including PTPRS, VCAN, and MXRA5, that operate within the stromal immune interface. These findings open new directions for understanding tumor immune regulation and highlight how reasoning based artificial intelligence can serve as a cognitive partner in precision immunotherapy research.

## Significance

This work presents a reasoning centered, multi LLM approach that turns biomarker discovery into an interpretable, evidence driven process. Using structured prompts, five run averaging per model, and cross model consensus across GPT 4o, GPT 5, and Gemini 2.5 Pro, we evaluated 478 Ig domain surfaceome genes from GSE245708 with 84 prostate cancer patients on a three point scale for immunotherapy, inflammation, and NF kappa B relevance. Ranking by mean scores identified a shared top ten set, within which PTPRS, VCAN, and MXRA5 emerged as novel, biologically plausible regulators lacking active drug programs. By linking model reasoning to literature evidence, pathway context, and extracellular tractability, this framework provides a reproducible path from RNA sequencing data to actionable immunotherapy targets.

## Key Points

- A structured, multi LLM reasoning workflow evaluates surfaceome genes with a three point scoring scheme and five independent runs per model, followed by cross model averaging for robust prioritization.
- The study analyzes GSE245708 with 84 patients, filters to 478 Ig domain surfaceome genes, and ranks all candidates by mean relevance across immunotherapy, inflammation, and NF kappa B categories.
- The common top ten set includes known regulators such as CD276 and B2M, and highlights PTPRS, VCAN, and MXRA5 as previously unrecognized stromal immune checkpoint like candidates.
- Only F3 has an FDA approved therapy, underscoring a translational gap and motivating target development for high scoring but untreated candidates.
- Next steps include mapping extracellular binding partners and defining how these interactions influence immune cell activation, polarization, and infiltration, to guide rational antibody or biologic design.

## Author contributions

N.P. and J.H.L. jointly conceptualized and designed the study. N.P. developed the IMMUNIA framework and performed computational analyses. J.H.L. curated and preprocessed the transcriptomic dataset and contributed to data interpretation and manuscript structuring. Both authors reviewed and approved the final version of the manuscript.

## Acknowledgement

Not applicable.

